# SolXplain: An Explainable Sequence-Based Protein Solubility Predictor

**DOI:** 10.1101/651067

**Authors:** Raghvendra Mall

## Abstract

**Motivation:** Protein solubility is a property associated with protein expression and is a critical determinant of the manufacturability of therapeutic proteins. It is thus imperative to design accurate *in-silico* sequence-based solubility predictors.

**Methods:** In this study, we propose SolXplain, an extreme gradient boosting machine based protein solubility predictor which achieves state-of-the-art performance using physio-chemical, sequence and novel structure derived features from protein sequences. Moreover, SolXplain has a unique attribute that it can provide explanation for the predicted class label for each test protein based on its corresponding feature values using SHapley Additive exPlanations (SHAP) method.

**Results:** Based on an independent test set, SolXplain outperformed other sequence-based methods by at least 2% in accuracy and 2% in Matthew’s correlation coefficient, with an overall accuracy of 78% and Matthew’s correlation coefficient of 0.56. Additionally, for fractions of exposed residues (FER) at various residual solvent accessibility (RSA) cutoffs, we observed higher fractions to associate positively with protein solubility, and tripeptide stretches that contain one isoleucine and one or more histidines, to associate negatively with solubility. The improved prediction accuracy of SolXplain enables it to predict protein solubility with greater consistency and screen for sequences with enhanced manufacturability.

## 1 Introduction

Protein solubility is an essential physiochemical property that determines the success of many biotechnical and scientific applications, including the production of therapeutic proteins and drugs based on antibodies. However, many proteins when expressed with a standard production procedure in *Escherichia coli*, have low solubility, which reduces their manufacturing capability. Enhancement of protein solubility in experimental settings is usually achieved through the use of low temperatures, weak promotors, modified growth media, or optimization of expression conditions [1, 2].

The primary determinant of protein solubility are the amino acid sequences in a protein. Previous studies showcased several correlations between amino acid sequence properties and protein solubility, such as the content of turn-forming and charged residues, the quantity of hydrophobic stretches, content of different types of residues, or the length of the protein sequence [3, 4, 5, 6].

This fact led to the development of many *in-silico* bioinformatics tools predicting the solubility propensity of proteins from their primary structure. Their aim is to replace the costly experimental procedures by pre-selecting the most promising protein sequences *in-silico*. Several protein solubility prediction tools include PROSO [7], SOLpro [2], CCSOL [8], PROSO II [9], PaRSnIP [10] and DeepSol [11]. Majority of these methods follow a two-step process: a) feature engineering and selection; b) protein solubility propensity prediction i.e. distinguishing soluble proteins from insoluble ones. PROSO tool uses SVM with Gaussian kernel and a Naive Bayes classifier to build its classifier. SOLpro utilizes a two-stage SVM with sequential minimal optimization to build the protein solubility predictor. CCSOL uses an SVM classifier and identifies coil/disorder, hydrophobicity, *β*-sheet and *α*-helix propensities as most discriminative features. PROSO II method uses a Parzen window model with modified Cauchy kernel and a two-level logistic classifier. PaRSnIP uses a gradient boosting machine algorithm which provides feature importance during training phase. Recently, a deep learning technique called DeepSol [11], showcased that by just using the raw protein sequences as input, it can perform as well as its competitors. Furthermore, by augmenting few additional phyiso-chemical features, it can easily surpass all other *in-silico* sequence-based protein solubility predictors. However, it is difficult to determine the biological relevance of the features engineered by deep learning as mentioned in [11, 12].

To overcome the limitations of exisiting methods, we propose SolXplain, an XGBoost based model [13] using several well-known physio-chemical and sequence derived features. Additionally, SolXplain uses several novel secondary structure, disorder, disulfide bonds related, torsion angles and contact number related features extracted from the SCRATCH suite [14], DISOPRED version 3.16 [15], DISpro [16] and SPIDER3 [17] respectively. XGBoost is an optimized version of gradient boosting machine [18], which has been shown to perform very well on several bioinformatics problems, such as gene regulatory network reconstruction [19, 20, 21], transmembrane protein solubility [22] etc. Moreover, SolXplain has a unique attribute that can provide explaination for the predicted class label for each test protein based on its corresponding feature values using SHapley Additive exPlanations (SHAP) [23] algorithm. Our primary contributions include:

1. Engineering of novel structure, disorder, disulfide bonds, torsion angles and contact number related features from the protein sequence using SCRATCH suite, DISOPRED, DISpro and SPIDER3 respectively.
2. Usage of an XGBoost model enables SolXplain to outperform existing methods for various evaluation metrics on an independent test set.
3. Provide explaination for SolXplain’s output by showing the most important features driving the model predictions towards soluble or insoluble class.

Figure 1 provides the steps undertaken by the proposed SolXplain model.

**Figure 1:**
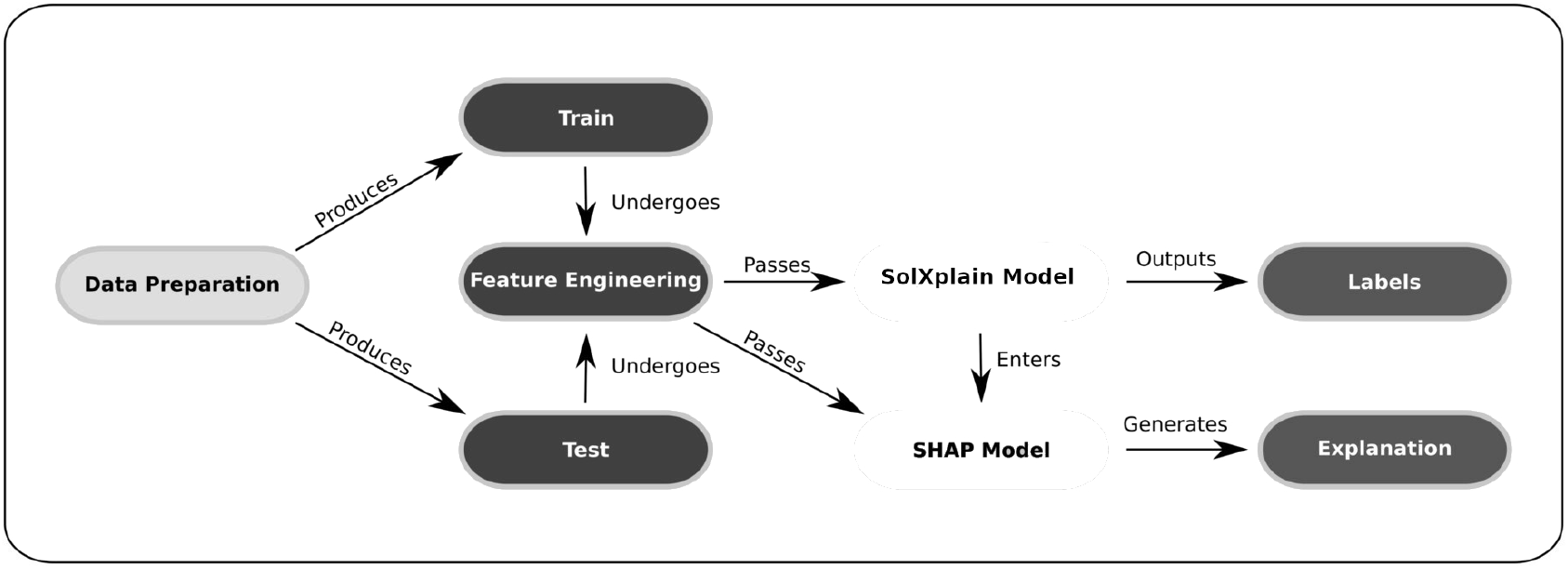
Flowchart followed by the proposed SolXplain model.

## 2 Materials and Methods

### 2.1 Overview

Our task is a binary classification problem i.e. distinguishing soluble proteins from insoluble ones. Our goal is to learn a function (*H*) which takes features engineered from a protein sequence i.e. **x** ∈ ℝ^*d*^ as input, and outputs a prediction score between [0, 1] ∈ ℝ i.e. *H*: **x** →[0, 1]. In this work, *H* is an XGBoost model [13], a white-box non-linear tree-based interpretable boosting machine that exploits the interactions between the engineered features.

### 2.2 Data Information

The original PROSO II dataset, consisting of 58, 689 soluble and 70, 954 insoluble sequences, was utilized as the training set. An independent test set of 1, 000 soluble and 1, 001 insoluble protein sequences first described by [24] was used as benchmark test set to evaluate the performance of SolXplain in comparison to other bioinformatic predictors.

We performed two main pre-processing steps to ensure sequence diversity within the training set and between training and independent test set. First, CD-HIT [25, 26] was used to reduce sequence redundancy within the training data set with a maximum sequence identity of 90%. Second, we excluded all training set sequences with a sequence identity of 30% or greater to any sequence in the independent test set to reduce the bias introduced by homologous sequences. The final training data set had 28, 972 soluble and 40, 448 insoluble sequences.

### 2.3 Feature Engineering

One of our primary contributions is to extract and design novel features which are useful for discriminating soluble proteins from insoluble ones. We devise three groups of features which are then used to train the SolXplain model (see Figure 2). The first set is composed of features based on global properties of the protein including sequence length (log(*L*)), molecular weight (log(Mol-Weight)), fraction of turn-forming residues, the average of hydropathicity (Gravy) and aliphatic indices, along with the total absolute charge. The second group of features are derived directly from the protein sequence and consist of frequencies of mono-(single amino acids, denoted AA), di-(two consecutive AAs) and tri-peptides (three consecutive AAs) within the protein sequences.

**Figure 2:**
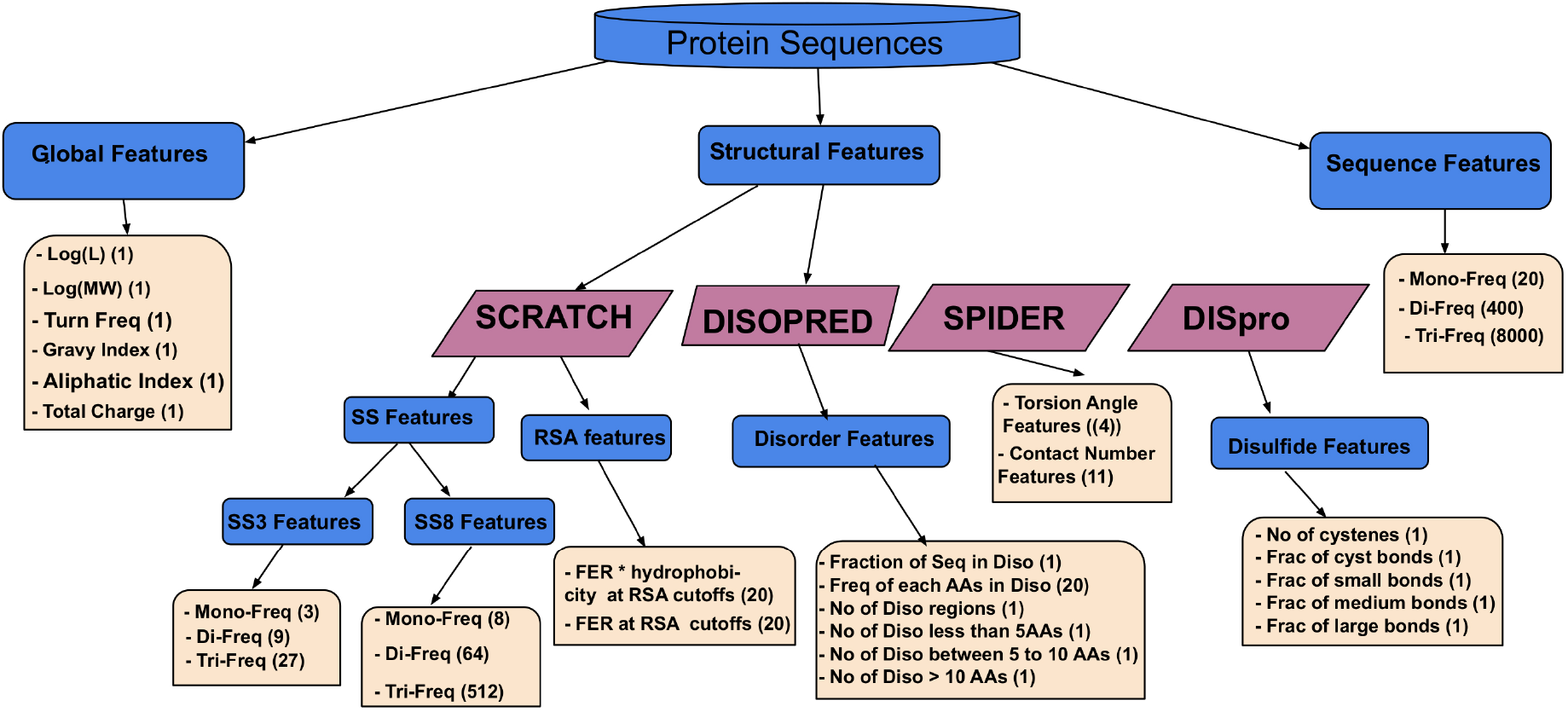
Different sets of features engineered for the SolXplain model. The number of features for each component is shown within parentheses.

The third group consists of structural features obtained from the protein sequence using SCRATCH [14], DISOPRED [15], DISpro [16] and SPIDER3 [17]. It has been shown previously [27] that SCRATCH based features are useful for protein fold prediction. We predict 3- and 8-state secondary structure (SS) information as well as the fraction of exposed residues (FER) at 20 different relative solvent accessibility (RSA) cutoffs (≥ 0%-≥95% cutoffs at 5% intervals). From the 3-state SS obtained via SCRATCH, we extract mono-(1 state i.e. turn, strand or coil), di-(two consective states) and tri-state (three consecutive states) frequencies for a given protein sequence. We follow a similar procedure for the more granular 8-state SS information as shown in Figure 2. Additionally, we multiply the FER by the average hydrophobicity of these exposed residues at different RSA cutoffs.

From DISOPRED, we obtain information about which AAs in the protein sequences are part of disordered regions as well as which AAs from the protein binding sites (PBS) of a protein are part of disordered regions. Given this information, we design features such as the fraction of the protein sequence which is disordered, frequency of each of the AAs (out of the 20 AAs) in disordered regions, number of disordered regions, number of disordered regions of length *<* 5 AAs, number of disordered regions of length between 5 and 10 AAs, and number of disordered regions of length *>* 10 AAs in a protein sequence. A similar set of features are extracted from the PBS of a protein.

From DISpro, we obtain information about cysteine disulfide bonds in the protein sequence. Since, the disulfide bonds are strong bonds, they are important for the structure and stability of proteins. We engineer features such as number of cysteine residues in protein sequence, ratio of number of cysteine bonds to ideal number of possible cysteine bonds, fraction of cysteine bonds between two residues which are less than 12 amino acids away from each other i.e. fraction of small bonds, fraction of cysteine bonds between two residues which are greater than 12 AA but less than 24 AA away from each other i.e. fraction of medium bonds and finally fraction of cysteine bonds between two residues which are greater than 24 AAs away from each other i.e. fraction of large bonds.

From SPIDER3, we extract features related to torsion angles (*ψ* and *φ*), half sphere exposure (HSE) and contact number (CN) which are important to determine the structure and stability of a protein. We engineer 4 features determining density of residues in *α*-helix, *β*-sheet, lefthand *α* and the remaining regions from the Ramachandran plot [28] based on the *ψ* and *φ* angles predicted by SPIDER3. Higher fraction of residues in *α*-helix, *β*-sheet and lefthand *α* regions of the Ramachandran plot are characteristics of a stable protein structure. We finally extract 11 features related to contact numbers and HSE of residues in a protein sequence such as average contact number, HSE value of residues in the range set *S* = {≤ 10, > 10 & < 20, ≥ 20}, AAs away from these residues.

In total, we include 9158 features for each protein sequence. In contrast to other sequence-based predictors, we don’t perform a feature selection step to exclude features, rather we rely on the XGBoost model to prioritize the most important features and filter out irrelevant ones.

### 2.4 Methods

#### 2.4.1 Gradient Boosting Machine

The SolXplain model is an optimized version of the white-box, non-linear, ensemble gradient boosting machine (GBM) [18, 29] called XGBoost [13]. Gradient Boosting is a machine learning method based on a constructive strategy by which the learning procedure will additively fit new models, typically decision trees and repetitively leverage the patterns in residuals to provide a more accurate estimate of the response variable (soluble versus insoluble proteins).

#### 2.4.2 XGBoost Algorithm

Tree boosting is a learning technique to improve the classification of weaker classifers by repeatedly adding new decision trees to the ensembles. XGBoost [13] is a scalable machine learning technique for tree boosting. It was shown in [13] that its performance is better than other boosting algorithms.

The main components of XGBoost algorithm are the objective function and its iterative solution. The objective function is intialized to describe the model’s performance. Given the training dataset, 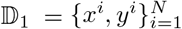 where *x*^*i*^ ∈ ℝ^*d*^, *d* = 9158, *y*^*i*^ ∈ ℝ and *N* denotes the number of total number of training samples. The predicted output 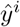 obtained from the ensemble model can be represented as: 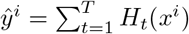, where *H*_*t*_(*x*^*i*^) represents the prediction score of the *t*^th^ decision tree for the *i*^th^ protein sequence in the training dataset. If the decision trees are allowed to grow unregulated, then the resulting model is bound to overfit [13]. Hence, the following objective has to be minimized:

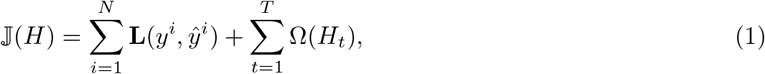

where **L** is the loss function and Ω(*⋅*) is the penalty that is used to prevent overfitting which is defined as: 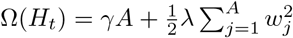, where *γ* and *λ* are the parameters which control the penalty for the number of leaf nodes (*A*) and leaf weights (*w*) respectively in the decision tree *H*_*t*_.

The objective function can be re-written as follows: 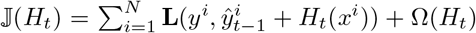. After applying a Taylor expansion and expanding Ω(*H*_*t*_), we obtain:

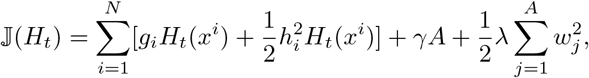

where 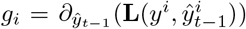 and 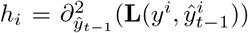 are the first and second order gradient statistics on the loss function. For a fixed tree structure *H*(**x**), where *I*_*j*_ = {*i*|*H*(*x*^i^) = *j*} is an instance of leaf node *j*, the optimal weight 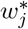 for leaf node *j* is:

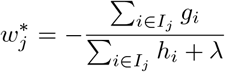

The corresponding optimal objective function then becomes:

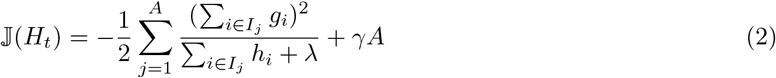

Equation 2 can be used as a scoring function to measure the quality of a tree structure *H*_*t*_ at the *t*^th^ iteration. This score is equivalent to the impurity score used for evaluating decision trees in random forests [30]. The authors in [13], devise a fast, greedy and iterative algorithm to identify these optimal tree structures.

### 2.5 Training

We train our XGBoost classifier on top of physio-chemical (global), sequence and structural features extracted from the protein sequence as mentioned earlier. Since our training set has slight imbalance, we weigh the samples belonging to soluble class by *α*, where 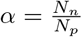 and *N*_*n*_ is the total number of insoluble proteins and *N*_*p*_ is the total number of soluble proteins in the training set.

The SolXplain classifier is based on several parameters, such as maximum depth of a tree (*M*), the learning rate (*ν*), the minimum child weight of a leaf node (*w*_*j*_), the sampling rate for features (*r*), and the regularization parameter *γ*. We keep the sampling ratio from training set parameter to a fixed value of 0.8. We then performed a hyper-parameter optimization procedure by varying these parameters over a grid of *M* × *ν* × *w*_*j*_ × *r* × *s* = 243 combinations. In particular with *M* ∈ {4, 7, 10,}, *ν* ∈ {0.01, 0.1, 0.2}, *w*_*j*_ ∈ {5, 10, 20}, *r* ∈ {0.1, 0.2, 0.3}, and *γ* ∈ {0.0, 0.25, 0.5}. We performed a ten-fold cross-validation for each of these combinations. Finally, we selected the XGBoost classifier which had the maximum ten-fold cross-validation area under the curve, which was obtained corresponding to the parameters *M* = 10, *ν* = 0.01, *w*_*j*_ = 20, *r* = 0.3 and *γ* = 0.0. The final SolXplain classifier had a maximum training accuracy of 85.93% and a maximum AUC of 0.94.

### 2.6 Evaluation Metrics

SolXplain method was compared against different in-silico sequence-based solubility predictors using several well-known metrics such as Accuracy (ACC), Matthew’s correlation-coefficient (MCC), Selectivity per class, Sensitivity per class and Gain per class. All these evaluation metrics are based on true positives (TP), true negatives (TN), false positives (FP), and false negatives (FN). The set TP represents the proteins which are soluble (class label 1) and for which SolXplain predicts *H*(**x**) ≥ 0.5. Similarly, the set TN consists of those proteins which are insoluble (class label 0) and for which SolXplain predicts *H*(**x**) *<* 0.5. The set FP represents those proteins whose true label is insoluble i.e. 0 but SolXplain predicts *H*(**x**) ≥ 0.5 and the set FN comprises proteins whose true label is soluble i.e. 1 but SolXplain estimates *H*(**x**) *<* 0.5. A detailed definition of these sets and importance of each of these evaluation metrics are provided in [11, 10, 31]. Higher the value of these metrics the better the performance of the predictor.

## 3 Experimental Results

### 3.1 Performance Comparison

The prediction performance of SolXplain was assessed using an independent test set first reported by [24]. We compared SolXplain to state-of-the-art solubility predictors DeepSol S2, DeepSol S1, PaRSnIP and PROSO II. We also implemented an optimized deep feed-forward neural network (DNNSol) on top of 9, 158 features for comprehensive comparison as depicted in Figure 3. SolXplain yielded a prediction accuracy of 77.79% and an MCC of 0.560, outperforming the state-of-the-art method DeepSol S2 by at least than 2% in accuracy and 2% in MCC (see Table 1).

**Table 1:** SolXplain outperforms all other predictors in 5 out of 8 evaluation metrics. Best results are highlighted in bold.

| Method | Accuracy | MCC | Selectivity (Soluble) | Selectivity (Insoluble) | Sensitivity (Soluble) | Sensitivity (Insoluble) | Gain (Soluble) | Gain (Insoluble) |
| --- | --- | --- | --- | --- | --- | --- | --- | --- |
| PROSO II | 0.672 | 0.344 | 0.667 | 0.678 | 0.687 | 0.657 | 1.335 | 1.355 |
| PaRSnIP | 0.740 | 0.481 | 0.758 | 0.725 | 0.705 | 0.775 | 1.517 | 1.448 |
| DeepSol S1 | 0.728 | 0.458 | 0.748 | 0.712 | 0.689 | 0.768 | 1.495 | 1.424 |
| DeepSol S2 | 0.766 | 0.546 | **0.843** | 0.718 | 0.654 | **0.878** | **1.686** | 1.434 |
| DNNSol | 0.764 | 0.531 | 0.800 | 0.735 | 0.702 | 0.825 | 1.602 | 1.469 |
| SolXplain | **0.780** | **0.560** | 0.773 | **0.786** | **0.791** | 0.786 | 1.547 | **1.572** |

**Figure 3:**
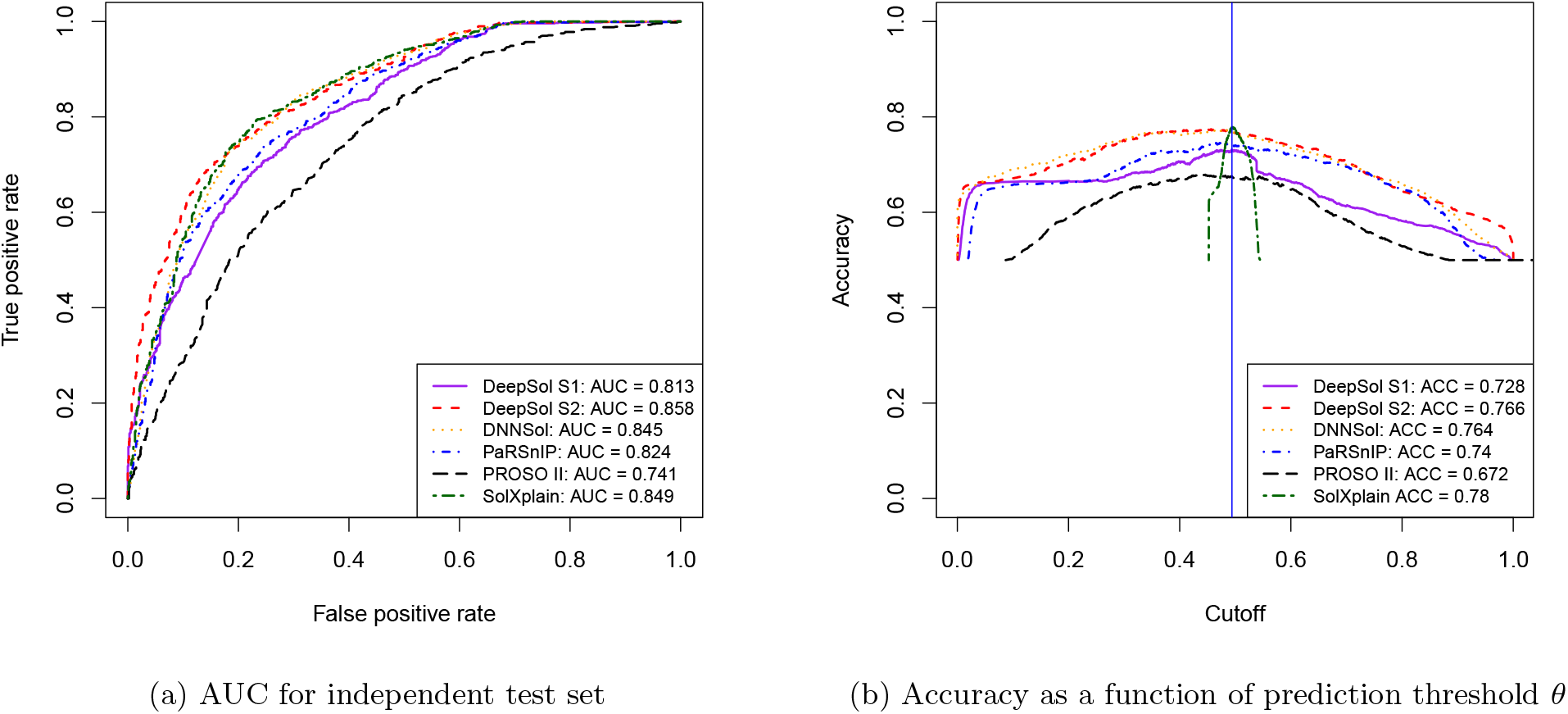
SolXplain performance comparison with other state-of-the-art solubility predictors.

SolXplain achieved stable values between 0.77 and 0.79 in sensitivity and selectivity metrics for both soluble and insoluble instances, whereas other predictors were mostly class biased. For example, the nearest competitors DeepSol S2 and DNNSol were better than SolXplain in detecting insoluble proteins as they attained high sensitivity for insoluble proteins and high selectivity for soluble proteins i.e. fewer insoluble proteins were misclassified as soluble proteins. However, DeepSol S2 and DNNSol were extremely poor, ≈ 15% and ≈ 9% worse than SolXplain in sensitivity (soluble class) respectively i.e. accurately identifying soluble proteins. Similarly, other classifiers such as DeepSol S1 and PaRSnIP were biased towards the insoluble class. However, their performance was worse than SolXplain for all evaluation metrics. From Figure 3b, we observed that the optimal cut-off (*θ*) to categorize the prediction of each predictor to soluble or insoluble class was *≈* 0.5.

By using score distribution plots as depicted in Figure 4, we were able to compare DeepSol S2 with DNNSol and our proposed approach SolXplain. It can empirically be illustrated that the score distributions for the 3 methods did not follow a normal distribution, since the violin plots don’t mimic a gaussian distribution. Hence, we used the Mann-Whitney-Wilcox (MWW) test [32] to compare pairwise score distribution for each class.

**Figure 4:**
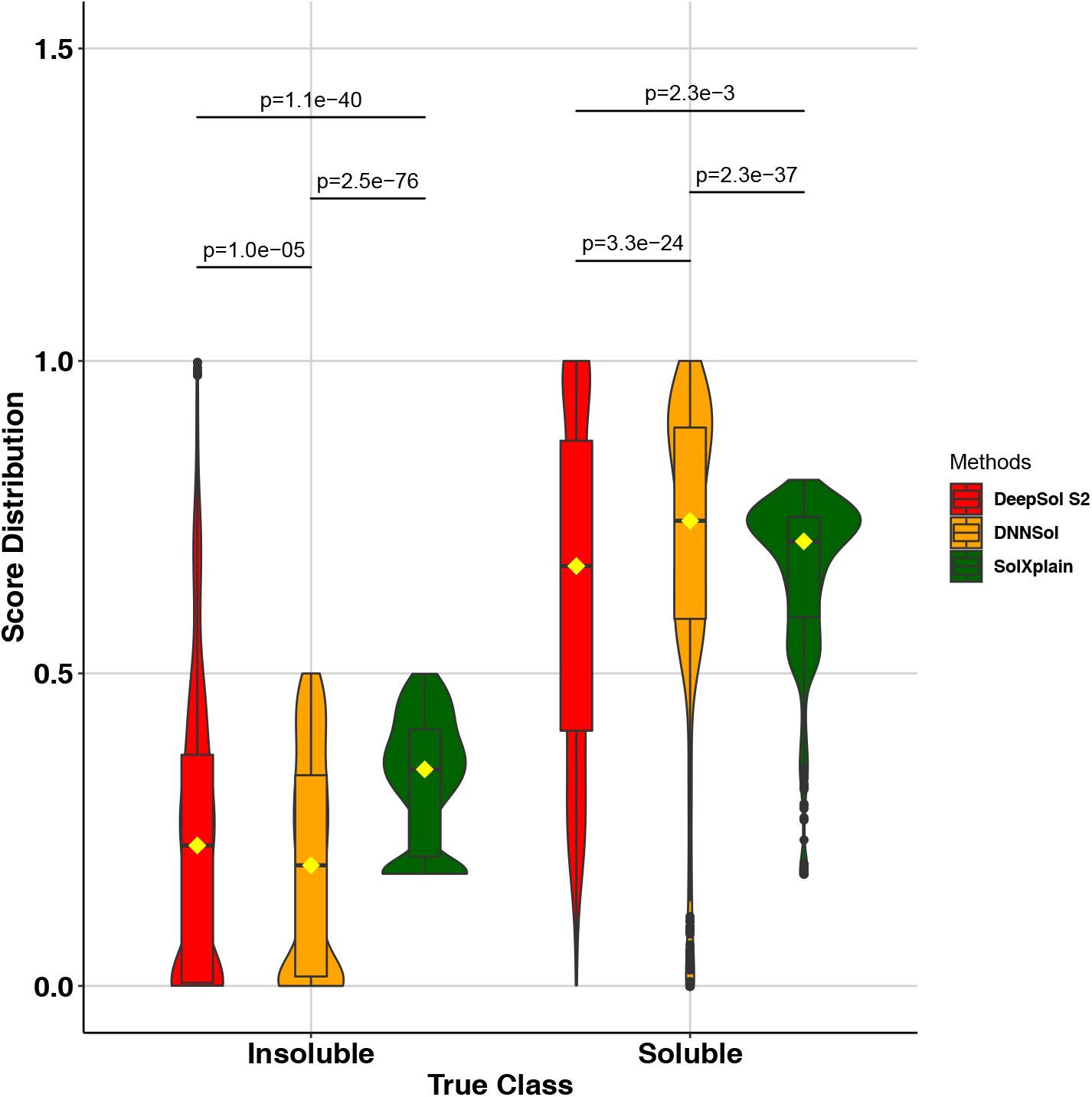
Comparison of score distribution of DeepSol S2, DNNSol with SolXplain. All the three solubility predictors have very different score distributions for class 0 or the insoluble class. However, the output score distribution of DeepSol S2 is not significantly different from SolXplain for the soluble class (*p* = 0.28). Here ‘p’ represents p-values and ‘yellow’ diamond corresponds to the mean of each score distribution.

We found that the score distributions of DeepSol S2 vs. DNNSol, DNNSol vs. SolXplain and DeepSol S2 vs. SolXplain were all different (pvalue *<* 1*e*-4) in the case of the insoluble class and the difference between the mean scores for DeepSol S2 vs. DNNSol and DNNSol vs. SolXplain were also statistically significant (pvalue *<* 1*e*-3) for the soluble class. Additionally, from the violin plot, we observed that the score distributions of DeepSol S2 and DNNSol have nearly similar shapes, indicative of both belonging to the deep learning class of machine learning methods, but very different from the density distribution achieved by SolXplain. This indicated that the underlying mechanism being used by SolXplain for identifying insoluble proteins was disparate from that being used by DeepSol S2 and DNNSol. However, the difference between the mean score for DeepSol S2 vs. SolXplain was not significantly different (pvalue= 0.28) even though their density distribution were quite dissimilar.

### 3.2 Feature Importance via XGBoost

An advantage of tree-based non-linear machine learning techniques, in contrast to black-box modelling techniques like support vector machines [33] and artificial neural networks [34], is that we can easily obtain feature/variable importance scores for all input features. The importance of a feature is the sum of information gained when splits (tree branching) are performed using that variable. A distinct benefit of using an XGBoost classifier is that out of all the 9158 features used during training, variables which are not used for optimal tree splits in the SolXplain model are pruned automatically and get a feature importance score of 0. We observe from Table 2 that only 499 features have non-zero feature importance scores.

**Table 2:** Variable importance percentages grouped by feature classes for the SolXplain model highlighting the contributions of features as depicted in Figure 2. Here total features (*>* 0) represents the number of variables from a feature class with *>* 0 variable importance scores.

| Feature Class | Total Features | Total Features (> 0) | Total Imp (%) |
| --- | --- | --- | --- |
| Log(L) | 1 | 1 | 0.18 |
| Log(MW) | 1 | 1 | 0.18 |
| Turn Freq | 1 | 1 | 0.06 |
| Gravy Index | 1 | 0 | 0.00 |
| Aliphatic Index | 1 | 1 | 0.09 |
| Total Charge | 1 | 1 | 0.10 |
| Mono Freq | 20 | 19 | 2.05 |
| Di Freq | 400 | 200 | 14.20 |
| Tri Freq | 8000 | 72 | 36.62 |
| Mono Freq SS3 | 3 | 3 | 0.24 |
| Di Freq SS3 | 9 | 8 | 0.57 |
| Tri Freq SS3 | 27 | 15 | 0.91 |
| Mono Freq SS8 | 8 | 7 | 0.86 |
| Di Freq SS8 | 64 | 22 | 2.13 |
| Tri Freq SS8 | 512 | 69 | 4.66 |
| FER at RSA cutoffs | 20 | 18 | 20.17 |
| FER at RSA cutoff × HP | 20 | 17 | 11.71 |
| Disorder Features | 25 | 17 | 1.95 |
| PBS in Disorder Features | 25 | 10 | 1.86 |
| Disulfide Features | 5 | 4 | 0.33 |
| Torsion Angle Features | 4 | 3 | 0.13 |
| Contact Number Features | 10 | 10 | 1.01 |
| All Features | 9158 | 499 | 100 |

We then analyzed the total feature importance contribution of all features according to their feature types/classes as shown in Figures 2 and Table 2. At the highest level, we had three macro classes of features including global, sequence, and structure derived features contributing 0.61%, 52.86%, and 46.53% respectively in the overall variable importance scores. From Table 2, we can observe that the maximum feature importance is associated with tripeptide frequencies (36.62%) followed by FER at different RSA cutoffs (20.17%). It is noteworthy that out of the 8, 000 features associated with tripeptide frequencies only 72 have non-zero feature importance. Thus, majority of the non-essential features are automatically pruned by XGBoost model. Figure 5 showcases the difference between the feature values for these top 3 variables in case of soluble versus insoluble proteins.

**Figure 5:**
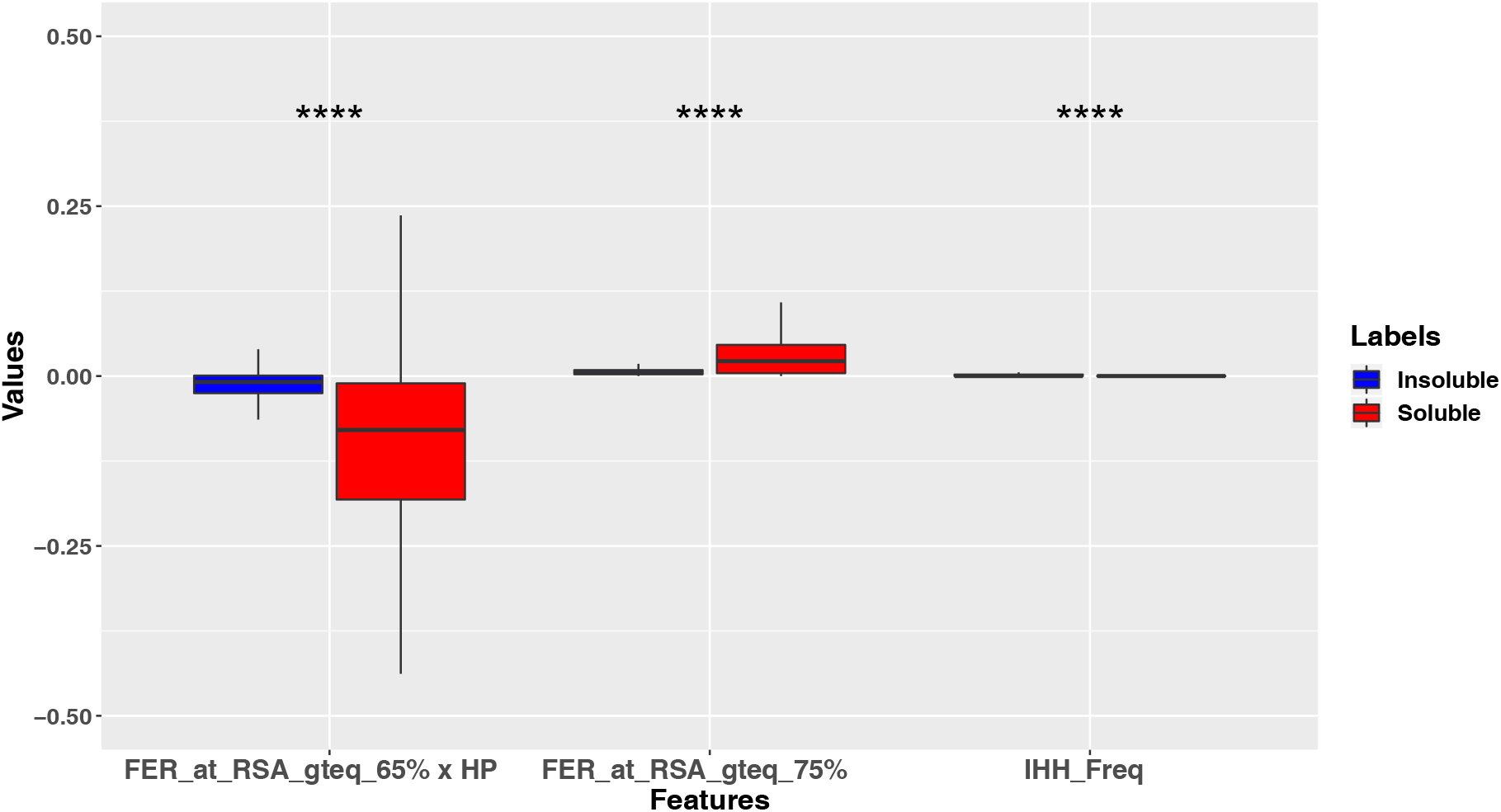
Distribution of the feature values of top 3 variables discriminating soluble from insoluble training protein sequences shown as box plots (****: P-value *<* 1*e*-4).

### 3.3 Model Explanation via SHAP method

A disadvantage of the inherent feature importance scores obtained from the XGBoost model is that the directionality is not apparent i.e. when a particular feature for a protein sample takes a high/low value, does it correspond to high/low feature importance. Moreover, at the test phase, it is not straightforward for white-box tree-based machine learning techniques to provide information about, say, the top *k* features driving the prediction to be soluble or insoluble class.

Recently, several techniques such as the LIME [35] and SHAP [23] methods have been proposed to overcome the aforementioned limitations. These methods have the ability to intepret feature importance scores from complex training models as well provide interpretable predictions for a test sample by grounding their reasoning on the top *k* features for that particular test instance. In our work, we use the SHAP (SHapley Additive exPlanations) method, a unified framework for intepretating predictions, as it was shown in [23] to outperform LIME and demonstrate that its predictions are better aligned with human intutions.

The SHAP method belongs to the class of additive feature attribution methods where a test instance prediction is composed as a linear function of features and satisfies 3 critical properties including local accuracy, missingness and consistency. The explicit SHAP regression values comes from a game-theory framework [36, 37] and can be computed as:

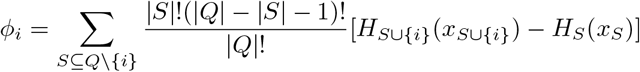

Here *Q* represents the set of all *d* features and *S* represents the subsets obtained from *Q* after removing the *i*^th^ feature and *φ*_*i*_ is an estimate of the importance of feature *i* in the model. In order to refrain from undergoing 2^*|Q|*^ differences to estimate *φ*_*i*_, the SHAP method approximates the Shapley value by either performing Shapley sampling [38] or quantitative input influence [39]. A detailed description of the SHAP method for model interpretation is available in [23].

We passed our SolXplain model along with the training set features to the SHAP method as shown in Figure 1 to obtain importance of features based on Shapley values. Figure 6 highlights the top 20 training features based on Shapley values. Moreover, it also provides directionality i.e. when a feature attains “high” or “low” values, the corresponding Shapley values are positive or negative. The positive Shapley values drive the predictions towards soluble class whereas the negative Shapley values influence the predictions to move towards the insoluble class. From Figure 6, we can observe that when top features such as FER at RSA cutoffs ≥ 75%, ≥ 70%, and ≥ 60% take high values, the corresponding Shapley values are positive driving model prediction to soluble class, whereas when these features take low values (i.e. closer to 0), the corresponding Shapley values are negative. Similarly, Figure 6 illustrates that when top features such as tripeptides: IHH and KIH frequencies are high, the corresponding Shapley values are negative and driving the model prediction to the insoluble class.

**Figure 6:**
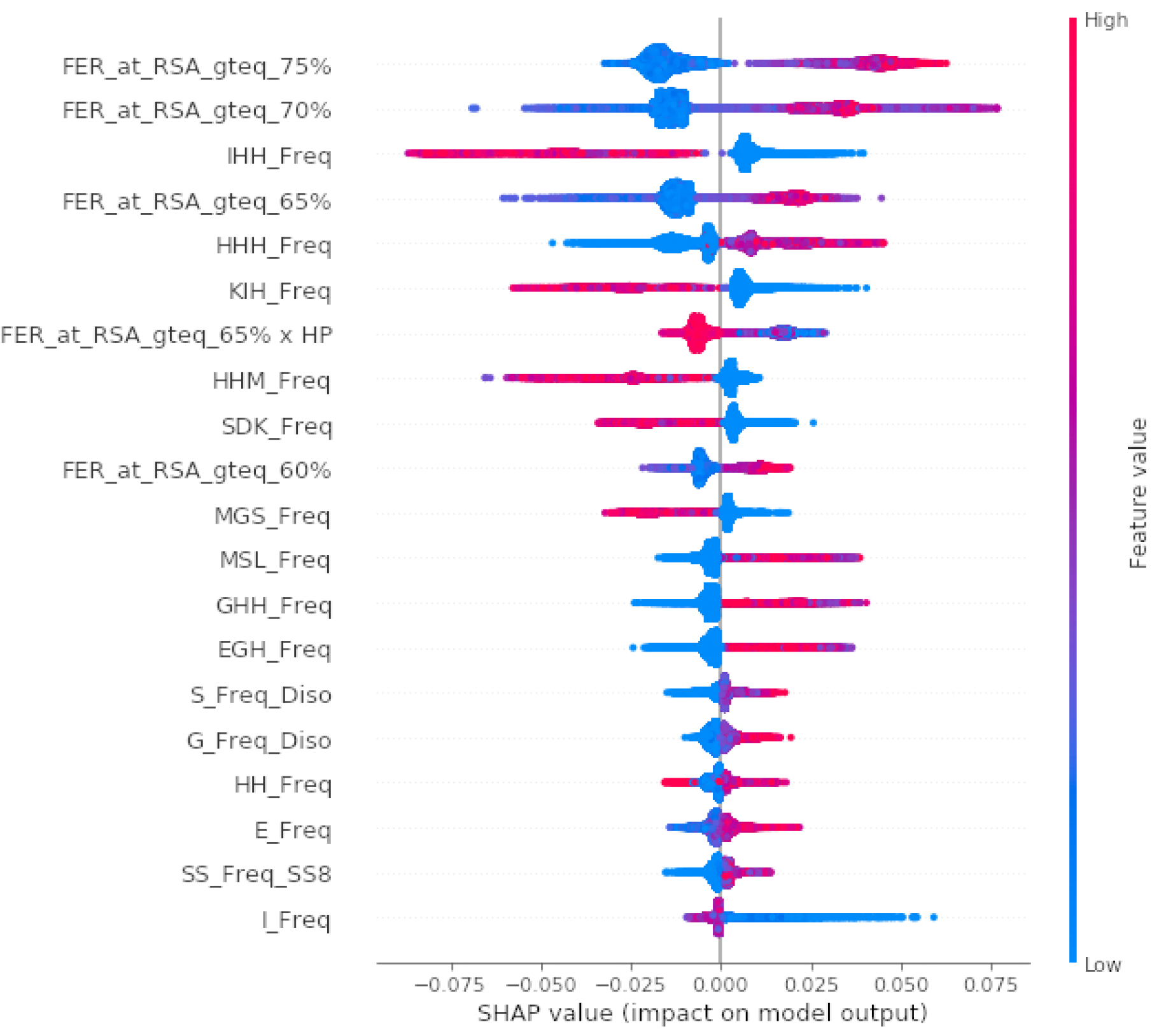
Top 20 features from SHAP [23] method. Here ‘gteq’ refers to *>*=.

### 3.4 Case Study

To better understand the varying SolXplain predictions, we analyzed the top 5 positive and top 5 negative features obtained via SHAP algorithm for two sample proteins. Acinetobacter baylyi pyrimidine nucleoside phosphorylase (3HQX) attained a relatively high SolXplain prediction score of 0.935, while the human AT-rich interactive domain-containing protein 3A’s (ARID3A) got a score of 0.725.

The first protein’s top 5 positive features were almost excusively made up of FER at different RSA cutoffs and had high positive Shapley values whereas the top 5 negative features had very small negative Shapley values. This drove the model prediction to be very close to 1 (0.935), suggesting the protein belonged to soluble class. We could observe from Figure 7 A and B, that there were several sequence of residues which were solvent accessible even at higher RSA cutoffs. For the second example protein again, the FER residues at different RSA cutoffs were the primary driver towards protein solubility prediction and attained high positive Shapley values. However, presence of tripeptide such as methionine-glycine-histidine and histidine-serine-histidine had a high negative Shapley value as depicted in Figure 7F. Thus, SolXplain achieved a relatively lower solubility score for ARID3A in comparison to 3QHX protein. Finally, from Figure 7 B and E, it was readily apparent that the phosphorylase had more RSA amino acids (depicted with light colors) than the second example protein ARID3A. Furthermore, ARID3A had 2 disordered regions at its terminal regions, which were not solved in the crystal structure, illustrated by dashed lines (7 D), that had antagonistic effect on the protein solubility.

**Figure 7:**
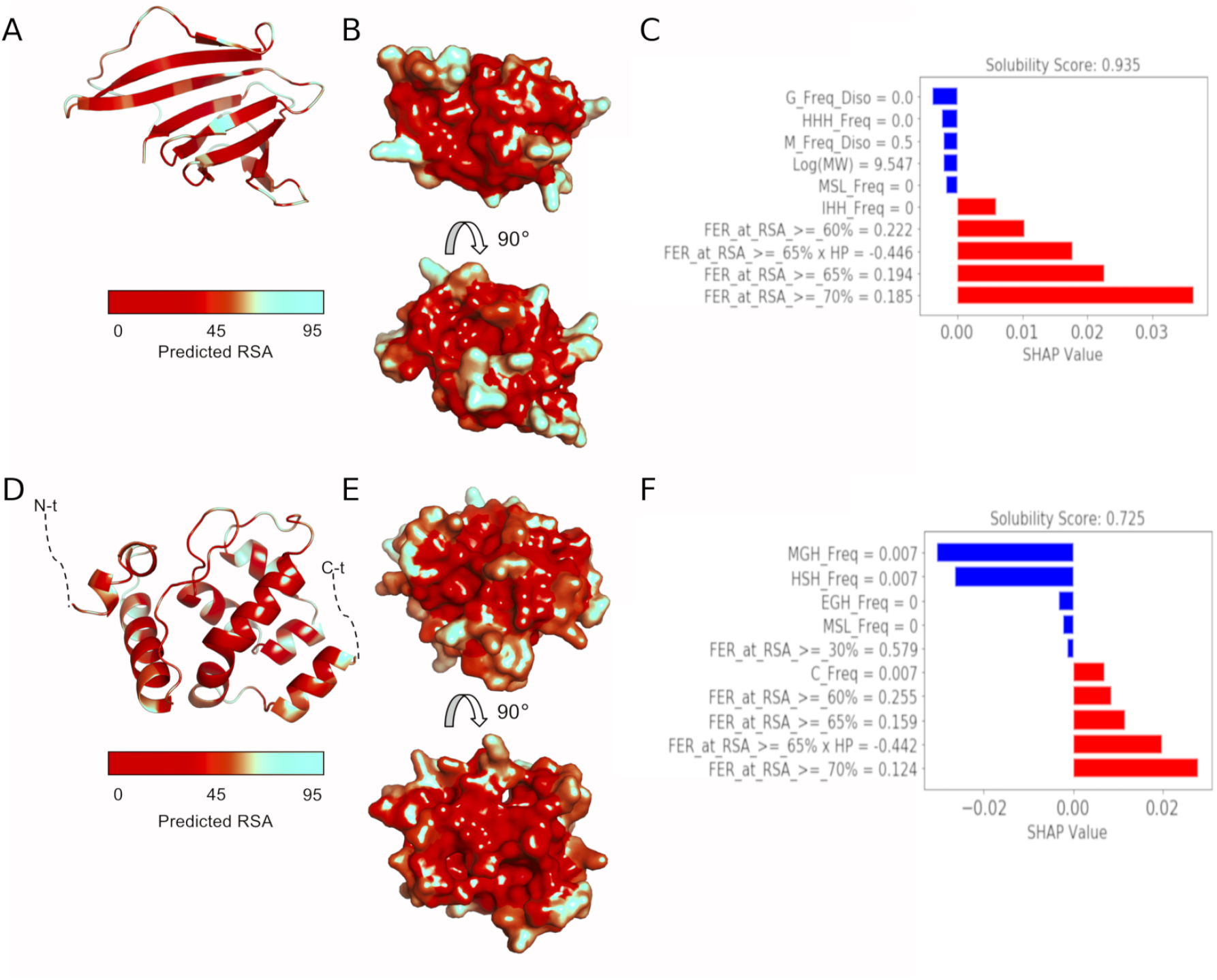
Top predictive features - relative solvent accessibilities and tripeptide frequencies - correlate with SolXplain scores. (A) Structure of pyrimidine/purine nucleoside phosphorylase with predicted RSA values mapped on structure in cartoon illustration (PDB ID: 3HQX, 108 residues). (B) Predicted RSA values shown on structure in surface representation. (C) Top 10 SHAP-features with corresponding Shapely values. (D) Structure of human AT-rich interactive domain-containing protein 3A (ARID3A, 145 residues) in cartoon illustration. (E) Predicted RSA values mapped onto structure an shown in surface representation. (F) Top 10 SHAP-features with corresponding Shapely values.

## 4 Discussion

The development of *in silico* sequence-based protein solubility prediction tools with high accuracy continues to be highly sought after. In this study, we introduce SolXplain, a solubility predictor that uses the XGBoost modeling technique and features that represent physio-chemical, sequence as well as structural properties of proteins. SolXplain outperforms, to the best of our knowledge, several existing sequence-based solubility predictors by *>* 2% in accuracy and *>* 2% in MCC.

The superiority of SolXplain over other predictors is due to three factors. The first factor is the choice of the machine learning method XGBoost. The non-linear optimized gradient boosting technique, XGBoost, is able to capture non-linear relationships between the features and the dependent vector, which makes its performance comparable to non-linear methods like SVMs [40]. Additionally, XGBoost reduces the bias of the model without increasing the variance, leading to better generalization performance. In addition, XGBoost has the ability to provide variable importance, making the model interpretable, which is a drawback of black-box methods like SVMs and deep learning [41]. The second factor is the choice of features. We include several features that provide information about the physio-chemical, sequence and structural properties of the protein of interest. Previous tools such as DeepSF [27] have shown that features extracted from SCRATCH suite are very helpful in correct protein fold recognition. We observe that the FER at different RSA cutoffs, average hydrophobicity of such residues and tripeptide frequencies play a very vital role in protein solubility prediction. An inherent advantage of XGBoost model is that it performs regularization i.e. feature pruning automatically, reducing the risk of overfitting and including only those features which help in discriminating the positive class from the insoluble ones. Thus, it has an advantage over two-stage methods like PROSO II and Solpro, which are susceptible to loss of information by explictly performing feature selection. Finally, unlike other sequence-based predictors, SolXplain has the ability to provide a meaningful explaination for each test sample using the SHAP method. This empowers biologists to quickly screen for good solubility targets and to attempt mutations of initial targets for soluble protein production reliably and with more information to reason about the effects of proposed modifications.

From the SolXplain model, (see Figure 5), we observe that the features with the highest variable importance were FER at RSA cutoffs 75%, FER at RSA cutoffs 65%× by average hydrophobicity of these residues and tripeptide IHH frequency. We notice from the training set that the FERs for the soluble set is significantly higher than the FERs for the insoluble set (see Figure 5, P-value *<* 1*e*-4), which is the reason that the FER is a dominant feature of the classifier. Additionally, from Figure 6, we also detect that the insoluble proteins tend to have higher frequency of tripeptides containing one isoleucine and one or more histidines in the protein sequence wheras soluble proteins have higher frequency of tripeptides containing only histidines. Interestingly, positively charged surface residues and polyhistidine-tags have been previously [42] correlated with protein solubility.

In this work, we develop SolXplain, a novel sequence-based protein solubility predictor that uses XGBoost technique and a large set of features depicting physio-chemical, sequence and structural properties of proteins. SolXplain not only outperforms all existing sequence-based protein solubility predictors, but is the first approach that provides meaningful interpretations for its prediction by using the SHAP method. SolXplain has relatively high sensitivity for soluble class (0.791) suggesting that it can efficiently select soluble proteins from an intial set of candidates, thereby, reducing the high attrition rate and the production cost.

